# Reduced histone gene copy number disrupts *Drosophila* Polycomb function

**DOI:** 10.1101/2023.03.28.534544

**Authors:** Jeanne-Marie E. McPherson, Lucy C. Grossmann, Robin L. Armstrong, Esther Kwon, Harmony R. Salzler, A. Gregory Matera, Daniel J. McKay, Robert J. Duronio

## Abstract

The chromatin of animal cells contains two types of histones: canonical histones that are expressed during S phase of the cell cycle to package the newly replicated genome, and variant histones with specialized functions that are expressed throughout the cell cycle and in non-proliferating cells. Determining whether and how canonical and variant histones cooperate to regulate genome function is integral to understanding how chromatin-based processes affect normal and pathological development. Here, we demonstrate that variant histone H3.3 is essential for *Drosophila* development only when canonical histone gene copy number is reduced, suggesting that coordination between canonical *H3.2* and variant *H3.3* expression is necessary to provide sufficient H3 protein for normal genome function. To identify genes that depend upon, or are involved in, this coordinate regulation we screened for heterozygous chromosome 3 deficiencies that impair development of flies bearing reduced *H3.2* and *H3.3* gene copy number. We identified two regions of chromosome 3 that conferred this phenotype, one of which contains the *Polycomb* gene, which is necessary for establishing domains of facultative chromatin that repress master regulator genes during development. We further found that reduction in *Polycomb* dosage decreases viability of animals with no *H3.3* gene copies. Moreover, heterozygous *Polycomb* mutations result in de-repression of the Polycomb target gene *Ubx* and cause ectopic sex combs when either canonical or variant *H3* gene copy number is also reduced. We conclude that Polycomb-mediated facultative heterochromatin function is compromised when canonical and variant *H3* gene copy number falls below a critical threshold.

## Introduction

To control access to information encoded in the genome, eukaryotes organize their DNA into chromatin, which regulates all DNA-dependent processes including transcription, DNA replication, and DNA damage repair (Allis CD 2007; Kornberg and Lorch 2020). The fundamental unit of chromatin is a nucleosome composed of approximately 150 bp of DNA wrapped around a histone octamer containing two copies of each of the four core histones: H2A, H2B, H3, and H4 (Luger et al. 1997). Tight control over histone levels is essential for normal genome function. For instance, mutations in *abo* and *mute*—which negatively regulate histone mRNA levels—reduce viability in *Drosophila melanogaster* (Berloco et al. 2001; Bulchand et al. 2010). In budding yeast, mutations that cause an accumulation of excess histone proteins result in impaired growth, DNA damage sensitivity, and chromosome loss (Meeks-Wagner and Hartwell 1986; Gunjan and Verreault 2003). Conversely, conditional repression of histone transcription during S phase impairs DNA replication and causes cell cycle arrest in yeast and fruit flies (Han et al. 1987; Sullivan et al. 2001; Gossett and Lieb 2012). Similarly, deletion of all *D. melanogaster* canonical histone genes leads to cell cycle arrest and embryonic lethality (Smith et al. 1993; Günesdogan et al. 2010; McKay et al. 2015). Histone chaperone mutations that reduce incorporation of histone proteins into chromatin cause spurious transcription, chromosome segregation defects, chromosomal rearrangements, and enhanced DNA damage (Clark-Adams et al. 1988; Nelson et al. 2002; Myung et al. 2003; Ye et al. 2003; Nashun et al. 2015; Mühlen et al. 2023a). For these reasons, precise regulation of histone mRNA and protein levels is critical for normal cell function and development, yet we have an incomplete understanding of the mechanisms involved.

Most research investigating the mechanisms of histone expression has focused on the canonical histone genes, which are synthesized in large amounts during S phase to properly package newly replicated DNA into chromatin. This work provides evidence supporting regulation at both the transcriptional and post-transcriptional levels (Marzluff and Duronio 2002; Duronio and Marzluff 2017). For example, in Chinese hamster ovary cells, canonical histone mRNA levels increase 35-fold as cells enter S phase (Harris et al. 1991). As cells exit S phase, canonical histone transcription is terminated and the corresponding mRNAs are rapidly degraded (Kaygun and Marzluff 2005; Eriksson et al. 2012). Coordinate expression among histone genes to maintain nucleosome subunit stoichiometry is also important; this requirement is reflected in the clustered arrangement and co-regulation of the canonical histone genes in multiple species, including *D. melanogaster,* yeast, and mammals (Lifton et al. 1977; Smith and Murray 1983; Eriksson et al. 2012). In the *D. melanogaster* histone gene complex (HisC, see **Figure 1A**), *H2A* and *H2B* share a bidirectional promoter, as do *H3* and *H4* (Lifton et al. 1977). Histone protein levels are also controlled post-translationally. For example, yeast histones that are not chromatin-bound are rapidly degraded, suggesting that excess histone proteins are deleterious to cell function (Singh et al. 2009). Moreover, during *Drosophila* oogenesis H2Av protein levels are regulated by Jabba, which binds H2Av and prevents degradation of excess histones (Stephenson et al. 2021).

**Figure 1.**
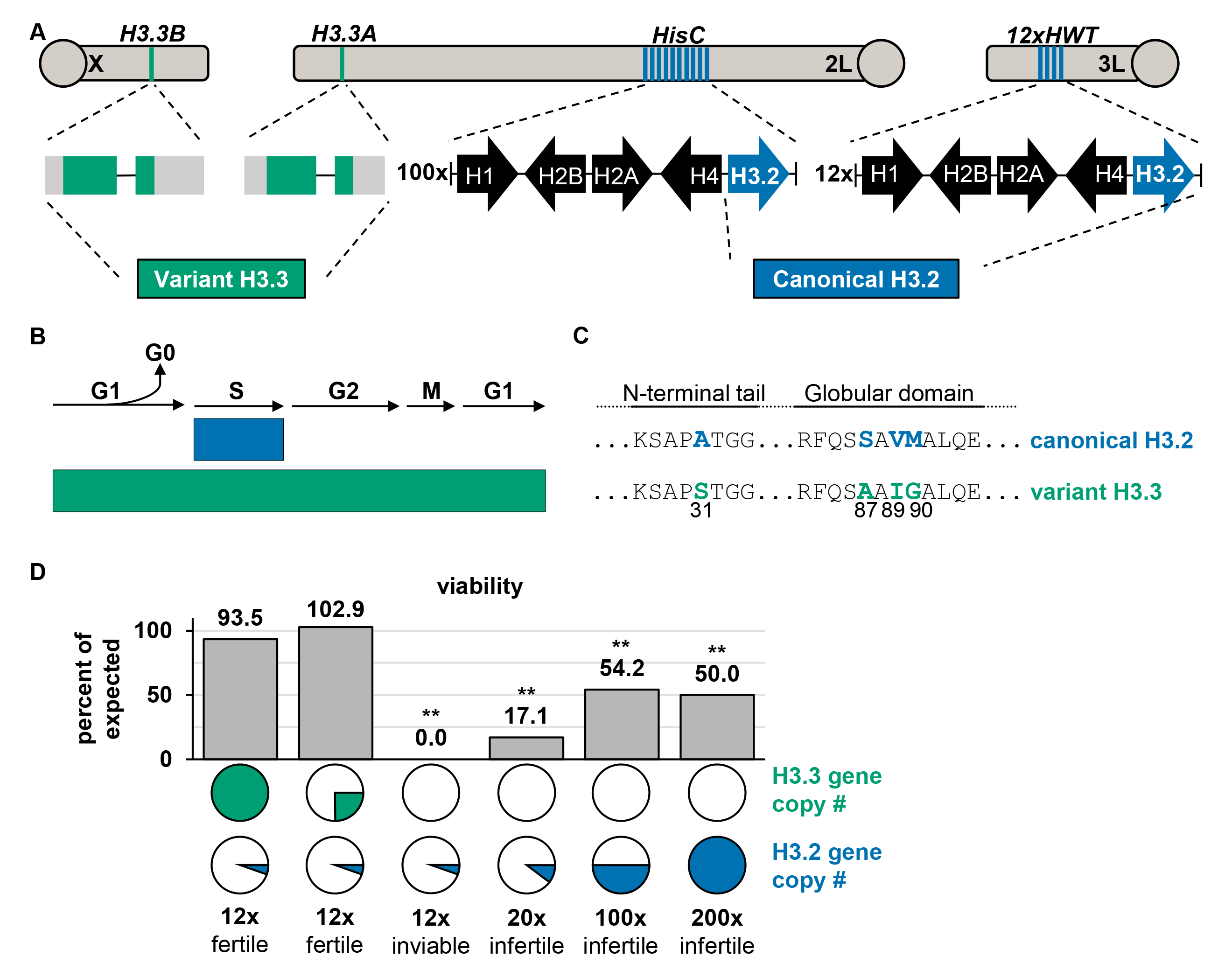
*H3.3* is required for viability when *H3.2* gene copy number is reduced. (A) Diagram of the genomic locations of the three H3 genes and the 12xHWT transgene. 100x and 12x indicate gene copy number. (B) Schematic model of *H3.2* and *H3.3* expression during the cell cycle. (C) Amino acid sequence differences of H3.2 and H3.3. (D) Bar plot of viability for the indicated genotypes. Circles represent the full complement of *H3.3* or *H3.2* gene copies present in the wild-type genome, and the color filling each circle (H3.3 green, H3.2 blue) indicates the number of gene copies present in each experimental genotype, e.g. a solid green circle indicates the 4 *H3.3* gene copies of the diploid *H3.3A* and *H3.3B* loci. The number of *H3.2* genes is shown below the blue circles, as well as whether surviving adults are fertile. Percent of expected genotypic frequencies based on Mendelian ratios. Asterisks indicate fewer than expected survive (chi-square test, ** p value < 0.01, see Figure 1 in File S1 for p values).

Whereas canonical histones are encoded by multiple genes that are expressed exclusively during S phase of the cell cycle (**Figure 1A, 1B**), an additional layer of complexity is provided by the expression of cell cycle independent histones (Franklin and Zweidler 1977; Verreault et al. 1996; Marzluff et al. 2002; Tagami et al. 2004). These so-called ‘variant’ histones are typically encoded by one or two genes and are expressed throughout the cell cycle (**Figure 1A, 1B**) (Urban and Zweidler 1983; Pantazis and Bonner 1984; Zweidler 1984; Brown et al. 1985; Piña and Suau 1987; Wunsch and Lough 1987; McKittrick et al. 2004; Tagami et al. 2004; Mito et al. 2005; Szenker et al. 2011; Maze et al. 2015; Tvardovskiy et al. 2017; Sauer et al. 2018). The tight control of canonical histone levels and the severe negative impact of histone mis-expression raises the possibility that coordinate regulation between canonical and variant histones is important for genome function and stability. For instance, in the early *Drosophila* embryo an increase in the ratio of variant H2Av to canonical H2A causes mitotic defects and reduces viability (Li et al. 2014). Here, we address the question of coordinate regulation between variant and canonical histone genes by focusing on those encoding histone H3.

In *D. melanogaster*, the non-centromeric H3 variant is encoded by two genes that produce identical proteins, *H3.3A* and *H3.3B*. (**Figure 1A**). Variant H3.3 differs from canonical H3.2 by only four amino acid residues, and these differences are highly conserved among other animals including humans (Malik and Henikoff 2003; Szenker et al. 2011) (**Figure 1C**). Three of the four residues are found in the globular domain and are known to modulate interactions with the histone chaperone complexes that deposit histones into chromatin (Grover et al. 2018). Canonical H3.2 is deposited during DNA replication by CAF-1 (chromatin assembly complex 1) (Smith and Stillman 1989; Verreault et al. 1996; Shibahara and Stillman 1999; Tagami et al. 2004; Sauer et al. 2018). Variant H3.3 is deposited into chromatin by the ATRX (alpha-thalassemia X-linked mental retardation protein) complex and the HIRA (histone cell cycle regulator) complex (Ahmad and Henikoff 2002; Tagami et al. 2004; Schneiderman et al. 2009; Goldberg et al. 2010; Lewis et al. 2010; Rai et al. 2011; Orsi et al. 2013; Ray-Gallet et al. 2018; Torné et al. 2020). Whereas H3.2 is deposited evenly genome-wide during replication, H3.3 is enriched at sites with high nucleosome turnover, including active regulatory elements, transcribed gene bodies, and pericentromeric regions (Ahmad and Henikoff 2002; McKittrick et al. 2004; Mito et al. 2005; Wirbelauer et al. 2005; Loyola and Almouzni 2007; Goldberg et al. 2010; Szenker et al. 2011; Martire and Banaszynski 2020 Jul 14). The fourth amino acid difference between H3.2 and H3.3 occurs at position 31 in the post-translationally modified N-terminal tail (Szenker et al. 2011). Position 31 is an alanine in H3.2 (H3.2A31) and a serine in H3.3 (H3.3S31), which can be phosphorylated (Hake et al. 2005; Armache et al. 2019; Martire et al. 2019; Sitbon et al. 2020). Other residues on the N-terminal tails of H3.2 and H3.3 are also differentially enriched in post-translational modifications (PTMs), likely due to their differential localization in the genome. Relative to H3.2, H3.3 is enriched with PTMs associated with active chromatin (e.g. H3K4me3) and depleted in marks associated with inactive chromatin (e.g. H3K9me2) (McKittrick et al. 2004; Hödl and Basler 2012). Although the mechanisms regulating canonical and variant histone mRNA and protein levels are distinct, we do not know if and how these mechanisms are coordinated to supply the necessary amount of each histone isotype across the genome. Here we use *D. melanogaster* to explore this question by examining the consequences of manipulating the relative number of canonical and variant *H3* genes.

Genetically manipulating histone gene copy number is challenging in many metazoans, including mice and humans, because canonical histones are encoded by multiple gene clusters located at distinct chromosomal locations (Marzluff et al. 2002b). *D. melanogaster* is a powerful organism to investigate the effects of altering histone gene copy number because all ∼100 haploid copies of the canonical histone genes are tandemly repeated (**Figure 1A**) and can be removed with a single genetic deletion, *ΔHisC* (Günesdogan et al. 2010). The ability to manipulate histone genes in *D. melanogaster* led to the discovery that canonical histone gene copy number is a modifier of position effect variegation (PEV), a genetic phenomenon associated with heterochromatin. Heterozygosity of the histone locus results in suppression of PEV, suggesting that histone abundance contributes to maintenance of epigenetic silencing of H3K9me3-marked constitutive heterochromatin (Moore et al. 1979; Moore et al. 1983; Sinclair et al. 1983). The ability to manipulate histone gene copy number in *D. melanogaster* has been extended in recent years. Replacement of all ∼200 copies (*200xWT*) of the canonical histone genes with a transgene containing 12 wild-type canonical histone gene repeat units (*12xHWT,* see **Figure 1A**) is sufficient to support development and provides a means of altering canonical histone gene copy number with precision (McKay et al. 2015).

Here, we report that *12xHWT* viability depends on expression of variant *H3.3* genes, whereas *200xWT* viability does not. This finding suggests that coordination of H3.2 and H3.3 protein levels is necessary for proper development when either *H3.2* or *H3.3* gene copy number is reduced. We conducted a screen to identify genes involved in the coordinated control of *H3.2* and *H3.3*. We identified a deficiency that uncovers *Yem*, a component of the HIRA histone chaperone complex, the function of which may be particularly important when *H3.2* gene copy number is reduced. Surprisingly, we also found that reduction of *Polycomb* (*Pc)* gene function decreases viability of flies that have reduced numbers of *H3.3* genes. Furthermore, we found that reductions in either *H3.2* or *H3.3* gene copy number disrupts Polycomb-mediated gene repression. Rather than *Pc* being involved in the coordinate expression of canonical and variant H3, we conclude from these findings that the appropriate balance of *H3.2* and *H3.3* genes is critical for the proper epigenetic silencing of developmental genes and maintenance of facultative heterochromatin function.

## Results

### *H3.3* is required for viability when *H3.2* gene copy number is reduced

To examine whether coordination between canonical *H3.2* and variant *H3.3* gene expression contributes to *Drosophila* development, we measured the effects of altering the relative number of canonical versus variant histone gene copies on viability and fertility. Zygotes lacking all canonical histone genes (*ΔHisC)* arrest early in embryonic development, and this lethality can be rescued with a transgene encoding 12 tandemly arrayed histone gene repeats (*12xHWT*), providing an opportunity to manipulate canonical histone gene dose over an ∼18-fold range (**Figure 1A, Figure 1D**) (McKay et al. 2015). Null mutations of either *H3.3A* or *H3.3B* have no effect on viability or fertility of flies containing the normal complement of canonical histone genes, but only 50% of the expected number of *H3.3A*, *H3.3B* double mutants (*H3.3^Δ^; 200xWT*) eclose as adult flies, which are infertile (**Figure 1D, Figure 1 in File S1**) (Sakai et al. 2009). *H3.3^Δ^* animals heterozygous for a *HisC* deletion (*H3.3^Δ^; 100xWT*) survive to adulthood at a similar frequency as *H3.3^Δ^; 200xWT* animals (54.2% and 50% of expected, respectively) (**Figure 1D**). However, reducing canonical histone gene copy number to 20 (*H3.3^Δ^; 20xWT*) results in only 17.1% of the expected number of adults (**Figure 1D**). A further reduction to 12 histone gene repeats (*H3.3^Δ^; 12xHWT*) results in a complete loss of viability of flies lacking variant *H3.3* genes (**Figure 1D**). *H3.3^Δ^; 12xHWT* lethality is rescued by one copy of either *H3.3A* or *H3.3B,* and the adults of these genotypes are fertile (**Figure 1D, Figure 1 in File S1**). Thus, we conclude that *H3.3* expression is necessary for completion of development when canonical histone gene copy number is reduced to 12, and that the probability of animals lacking *H3.3* to complete development increases with increasing numbers of canonical histone genes. Furthermore, our data support previous observations that *H3.3* is required for male and female fertility (**Figure 1D**) (Sakai et al. 2009). Collectively, these data suggest that *H3.3* compensates for reduced *H3.2* gene copy number to maintain a critical threshold of total H3 protein.

### H3.3 mRNA and protein levels do not change when *H3.2* gene copy number is reduced

Mechanisms that compensate for altered variant versus canonical histone genes could operate at many levels, including transcription, translation, histone deposition into chromatin, or histone protein turnover. For instance, increased *H3.3* expression could compensate for reduced *H3.2* gene copy number, potentially explaining why *12xHWT* animals do not survive in the absence of *H3.3* genes. We reasoned that measuring steady-state mRNA and protein levels could reveal evidence of such compensatory mechanisms. To determine whether *H3.3* steady-state mRNA levels are elevated in *12xHWT* animals, we compared *H3.3* mRNA levels in *12xHWT* versus *200xWT* control animals in an RNA-sequencing data set obtained from third instar larval brains. We found no significant difference in *H3.3A* or *H3.3B* mRNA levels in *12xHWT* compared to *200xWT* cells (**Figure 2A**). Consistent with these RNA-seq data, immunoblots of third instar larval wing imaginal discs show comparable levels of H3.3 protein in *12xHWT* and *200xWT* controls (**Figure 2B**). We conclude that compensation for reduced *H3.2* gene copy number in *12xHWT* animals does not occur via changes in the steady-state levels of *H3.3* mRNA or protein, as measured by RNA-sequencing or immunoblotting.

**Figure 2.**
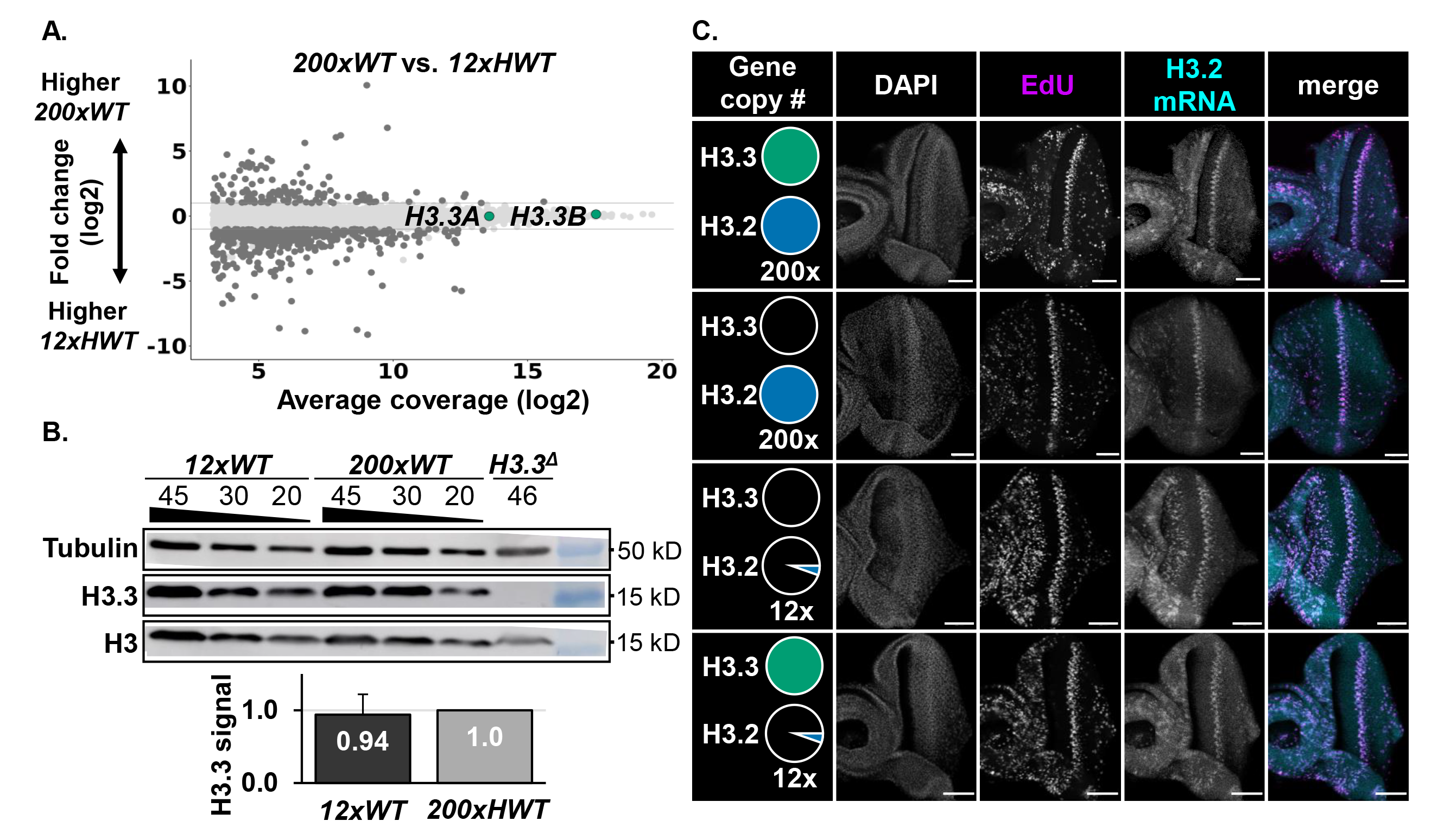
H3.3 mRNA and protein levels do not change when *H3.2* gene copy number is reduced. (A) MA plot showing fold change of normalized RNA-seq signal in wild-type (*200xWT*) vs. *12xHWT* for all transcripts (y-axis). Average coverage on the x- axis represents the mean expression level of a transcript. Differentially expressed genes are indicated in dark gray. Values for *H3.3A* and *H3.3B* transcripts are indicated (green). (B) Western blot of total H3, H3.3 and tubulin from *12xHWT* and *200xWT* wandering third instar larval wing imaginal discs. Dilution series with indicated number of wing discs for each genotype. H3.3^Δ^ third instar larvae were used as a negative control. Bar plot depicting the average fold change in H3.3 signal relative to *200xWT* signal normalized to tubulin. Error bars represent standard deviation of three biological replicates. (C) Confocal images of third instar imaginal eye discs stained for DAPI (grey), EdU (magenta), and H3.2 mRNA (cyan) for the four indicated genotypes, denoted as in Figure 1. The maximum projection of four adjacent slices is shown. Bars, 50 μM.

We considered the possibility that expression of *H3.2* becomes uncoupled from S phase upon loss of *H3.3*, thereby maintaining a pool of H3 outside of S phase even when *H3.3* genes are absent. To determine whether *H3.2* transcripts are present in cells outside of S phase in the absence of *H3.3* genes, we combined EdU staining with RNA-FISH in the developing eye of *200xWT*, *H3.3^Δ^, H3.3^Δ^; 12xHWT,* and *12xHWT* animals. We observed that *H3.2* mRNA is only detected in EdU positive cells in all four genotypes, indicating that *H3.2* transcription is not uncoupled from S phase when *H3.2* and/or *H3.3* gene copy number are reduced (**Figure 2C**).

### A genetic screen for genes sensitive to reduced histone H3 gene copy number

The inability to detect evidence of histone gene coordination at the molecular level motivated us to instead take an unbiased genetic approach. Performing a screen in a genotype with reduced variant and canonical histone gene copy number could potentially identify genes that (i) regulate histone gene expression, (ii) coordinate expression between variant and canonical histone genes, or (iii) are otherwise sensitive to reduced histone levels. As described above, *12xHWT* animals are viable and fertile at wild-type frequencies (**Figure 1D**); however, *H3.3^Δ^; 12xHWT* flies are inviable (**Figure 1D**). We therefore reasoned that we could identify other genes that when mutated would reduce the viability of *12xHWT* animals. Because having one copy of *H3.3A* or *H3.3B* is sufficient to retain viability in a *12xHWT* background (**Figure 1D**), we decided to screen using a *12xHWT* background that is further sensitized by the removal of both copies of *H3.3A*. We refer to this genotype as *H3.3A^Δ^;12xHWT* (**Figure 3A**).

**Figure 3.**
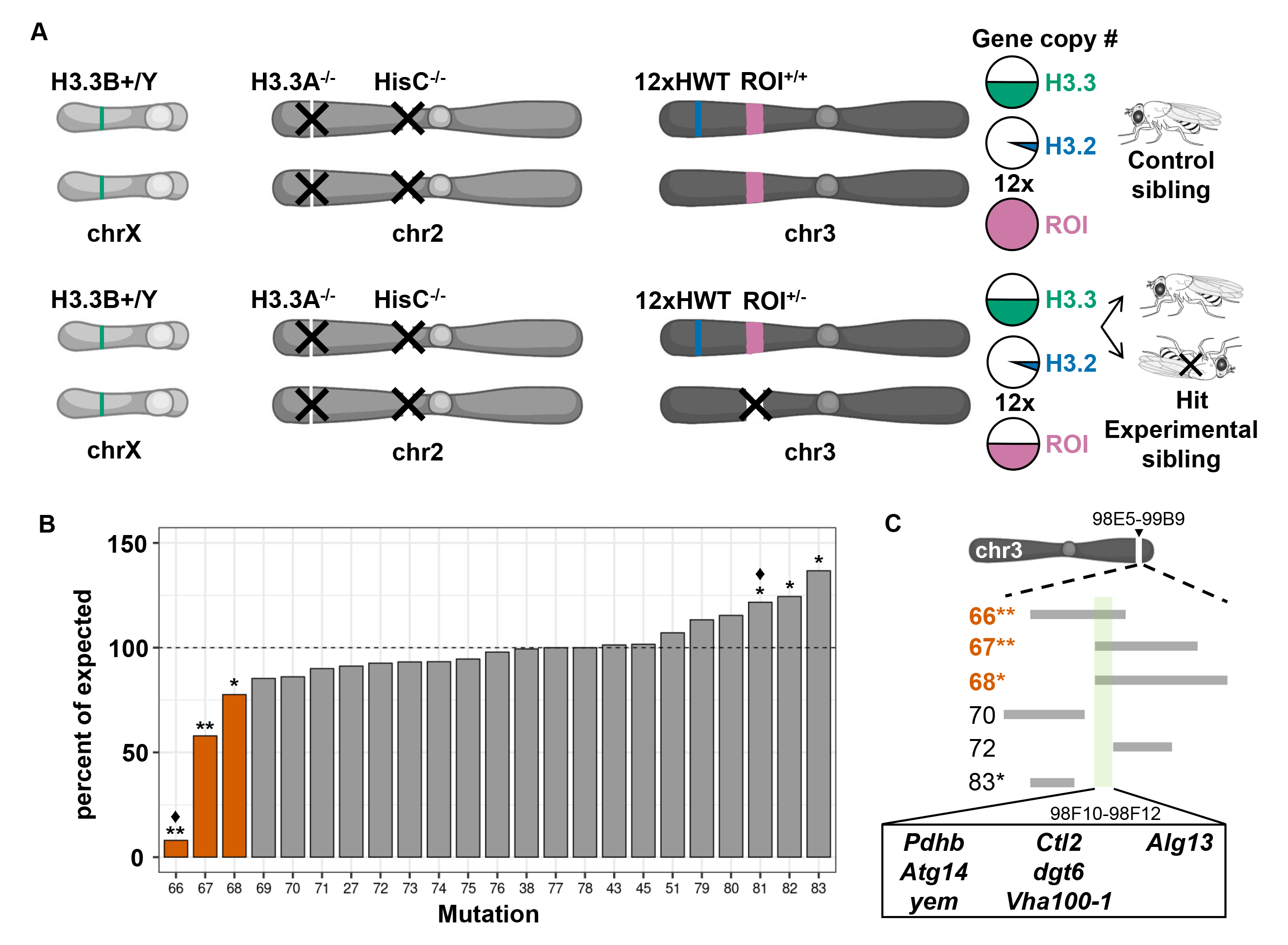
A genetic screen to identify genes required for survival when H3 gene copy is reduced. (A) Schematic of the screen used to identify regions of interest (ROI) on chromosome 3 that when heterozygous or hemizygous reduce adult viability of the *H3.3A^Δ^; 12xHWT* experimental genotype compared to control siblings. H3 gene copy number indicated as in Figure 1. Pink fill indicates gene copy number of the ROI. (B) Bar plot of viability for 23 mutations on chr3L and chr3R (see Table 1). Asterisks indicate statistical significance by Chi-square test (** p value < 0.01, * p value < 0.05, ♦ potential haploinsufficiency irrespective of *H3* gene copy number, see Figure 3 in File S1 for p values). Deficiencies covering region of interest indicated by orange bars. Data are plotted as the percent of *H3.3A^Δ^; 12xHWT* animals inheriting the deficiency chromosome relative to sibling animals inheriting the homologous balancer chromosome, which establish the expected percentage (dashed line). (C) Diagram of deficiencies that delineate a region of interest on chr3R from 98E5-99B9. The overlapping region specific to the positive hits from 98F10-98F12 contains seven genes (green bar).

**Table 1.** Chromosome 3R and 3L mutation alleles tested in proof-of-principle screen for changes in viability.

|  |  |  |  |  |
| --- | --- | --- | --- | --- |
| **Number<sup>a</sup>** | **Deficiency** | **Start coordinate<sup>b</sup>** | **End coordinate<sup>b</sup>** | **p value<sup>c</sup>** |
| 27 | *Df(3L)BSC289* | 1332329 | 1628100 | - |
| 45 | *Df(3L)ED4287* | 1795442 | 2551761 | - |
| 79 | *Df(3L)ED4284* | 1795442 | 1963552 | - |
| 80 | *Df(3L)BSC385* | 2259731 | 2417382 | - |
| 38 | *Df(3L)BSC730* | 12156077 | 12836424 | - |
| 51 | *Df(3L)ED4606* | 16080584 | 16773223 | - |
| 43 | *Df(3L)BSC220* | 18965662 | 19164368 | - |
| 77 | *Asf1[1]* | 19619559 | 19619559 | - |
| 78 | *E2F[rM729]* | 21626545 | 21626545 | - |
| 82 | *Df(3R)BSC527* | 22626930 | 23111808 | + |
| 71 | *Df(3R)ED6220* | 24543798 | 25183773 | - |
| 73 | *Df(3R)10-65* | 81F | 81F | - |
| 76 | *Df(3R)XNP[1]* | 25477868 | 25482834 | - |
| 81<sup>♦</sup> | *Df(3R)ro80b* | 97D1 | 97D13 | + |
| 75 | *Df(3R)BSC460* | 27937830 | 28461658 | - |
| 83 | *Df(3R)Exel6210* | 28674961 | 28991018 | + |
| 66<sup>♦</sup> | *Df(3R)BSC874* | 28675029 | 29191671 | ++ |
| 70 | *Df(3R)BSC789* | 28820134 | 29040507 | - |
| 67 | *Df(3R)BSC500* | 29112527 | 29675700 | ++ |
| 68 | *Df(3R)BSC501* | 29112527 | 29724685 | + |
| 72 | *Df(3R)ED6310* | 29138895 | 29512153 | - |
| 74 | *Stg[4]* | 29252826 | 29255800 | - |
| 69 | *Df(3R)BSC547* | 29621303 | 29821399 | - |
| <sup>a</sup> ♦ suggested haploinsufficiency |  |  |  |  |
| <sup>b</sup> Breakpoints are as determined by the Bloomington stock center. |  |  |  |  |
| <sup>c</sup> +, *p*-value < 0.05; ++, *p*-value < 0.01. *P*-values obtained from a Chi-square test. |  |  |  |  |

First, we conducted a proof of principle screen using single gene loss of function alleles or deficiencies covering genes with potential roles in histone function. We tested histone H3 chaperones (e.g. *Asf1*, *Yem, Xnp*), cell cycle regulators (e.g. *E2F*, *Stg*), and genes involved in the control of histone mRNA synthesis or chromatin regulation (e.g. *Slbp*, *wge*, *Arts, Dre4*). We performed crosses that produced *H3.3A^Δ^; 12xHWT* progeny heterozygous for individual mutations or deficiencies and determined whether the progeny had reduced or increased viability compared to *H3.3A^Δ^; 12xHWT* control siblings. Of the 23 mutations tested, heterozygosity of three deficiency mutations—*Df(3R)BSC874* (Df 66), *Df(3R)BSC500* (Df 67), and *Df(3R)BSC501* (Df 68)—resulted in a significant reduction in viability of *H3.3A^Δ^; 12xHWT* flies compared to control siblings, with only 8.4%, 57.9%, and 77.6% of expected surviving to adulthood, respectively (**Figure 3B, Table 1**). Interestingly, three other deficiencies Df(3R)ro80b (81), Df(3R)BSC527 (Df 82), and Df(3R)Exel6210 (Df 83) resulted in a significant increase in viability of *H3.3A^Δ^; 12xHWT* flies compared to control siblings, but we did not pursue these further (**Figure 3B, Table 1**). Notably, the three deficiencies that reduce *H3.3A^Δ^; 12xHWT* viability overlap the same 76.6 kb region on chromosome 3R (chr3R) (**Figure 3C**). To further define the genomic region responsible for the genetic interaction, we tested three other deficiency mutations that overlap this region. *Df(3R)BSC789* (Df 70), *Df(3R)ED6310* (Df 72), and Df 83, which overlap other regions of Df 66, Df 67, and Df 68, did not result in a significant reduction in viability. Therefore, the region of interest is limited to a 26.4 kb region defined by the left breakpoints of Df 67 and Df 72 at genomic positions 98F10 and 98F12 (**Figure 3C**). Seven annotated genes reside within this region of interest: *Alg13*, *Atg14*, *dgt6*, *Ctl2*, *Pdhb*, *Vha100-1*, and *yem*, an H3.3 specific chaperone (**Figure 3C**). Thus, one possible explanation for these genetic results is that heterozygosity of *yem* attenuates incorporation of H3.3 protein (derived from the *H3.3B* locus) into chromatin enough to reduce the viability of *H3.3A^Δ^; 12xHWT* flies. This targeted screen confirms that our genetic paradigm can identify mutant loci that when hemizygous cause sensitivity to a reduction in *H3.2* and *H3.3* gene copy number.

To expand our search for such loci, we screened the left arm of chromosome 3 (chr3L) using the Bloomington Stock Center Chr3L Deficiency Kit, which consists of 77 stocks that cover 97.1% of the chr3L euchromatic genome (**chr3L Df kit stocks in File S1**) (Cook et al. 2012; Roote and Russell 2012). Fourteen of the deficiency mutations were excluded from the screen because they carry a mini-white genetic marker, resulting in an eye color that precludes identifying all progeny classes (**chr3L Df kit stocks in File S1**). Three additional deficiency mutations were not scored because the crosses failed (**chr3L Df kit stocks in File S1**). Of the remaining 60 deficiency mutations, two resulted in an increase in viability when heterozygous in *H3.3A^Δ^;12xHWT* flies (**Fig 4A, Table 2**), which we did not pursue further. By contrast, heterozygosity of eleven deficiencies caused significant reductions in viability of *H3.3A^Δ^;12xHWT* flies (**Figure 4A, Table 2**). Four of these deficiencies (Df 1, 4, 8, and 15, **Figure 4A**) also caused a significant reduction in viability of siblings with one copy of *H3.3A* and 112 copies of the canonical histone genes (*H3.3A^+/-^; 112xHWT*), suggesting haploinsufficiency. We did not pursue these hits further. Interestingly, of the remaining hits *Df(3L)BSC435* (Df 2) and *Df(3L)BSC419* (Df 3) overlap the same 300 kb region of chr3L (**Figure 4B**). To map the genetic interaction in greater detail, we obtained two additional deficiencies—*Df(3L)BSC418* (Df 52) and *Df(3L)BSC836* (Df 36)—that overlap this same region of chr3L and observed no changes in viability of *H3.3A^Δ^;12xHWT* flies compared to control siblings (**Figure 4A-B**). Therefore, the genomic region spanning 77.76 kb on chr3L between cytological positions 78C6 and 78C8 impairs viability of flies with reduced *H3.3* and *H3* gene copy number.

**Figure 4.**
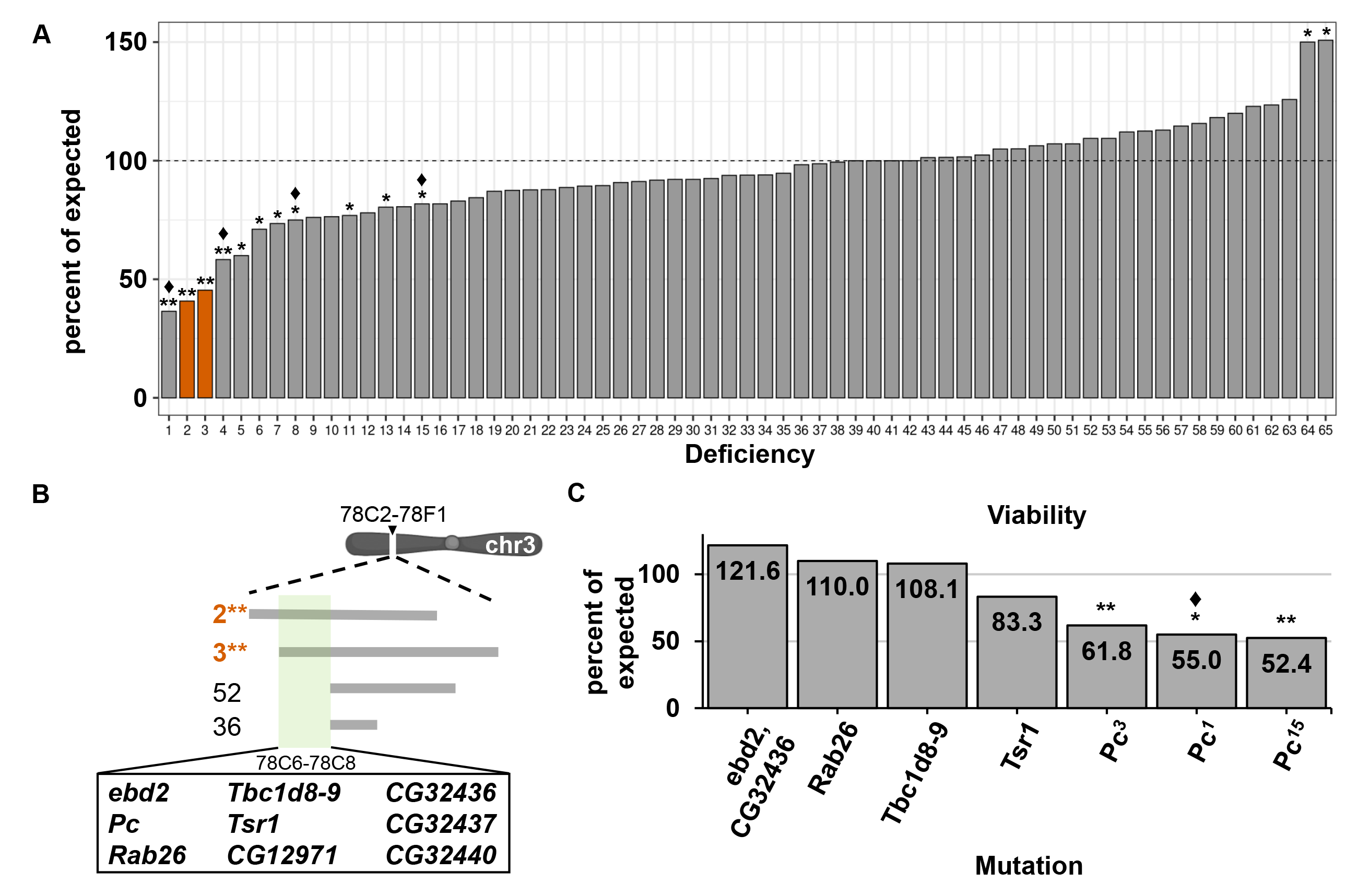
Screen of chromosome 3L deficiencies identifies histone gene copy number as a modifier of *Polycomb*. (A) Bar plot of viability for 65 chr3L deficiency mutations (see Table 2). Orange bars indicate two overlapping deficiencies that scored positive and uncover the *Pc* locus. Data are plotted as in Figure 3. (B) Diagram of deficiencies that delineate a region of interest on chr3L from 78C2-78F1. The overlapping region specific to the positive hits from 78C6-78C8 contains nine genes (green bar). *Df(3L)BSC435* (Df 2) and *Df(3L)BSC419* (Df 3) correspond to the orange bars in panel A. (C) Bar plot of viability of *H3.3A^Δ^; 12xHWT* animals heterozygous for the seven indicated mutations. Data are plotted as in Figure 3. Asterisks indicate statistical significance by Chi-square test (** p value < 0.01, * p value < 0.05, ♦ potential haploinsufficiency irrespective of *H3* gene copy number, see Figure 4 in File S1 for p values).

**Table 2.** Chromosome 3L deficiency alleles tested for changes in viability.

|  |  |  |  |  |
| --- | --- | --- | --- | --- |
| Number<sup>a</sup> | Deficiency | Start coordinate<sup>b</sup> | End coordinate<sup>b</sup> | p value<sup>c</sup> |
| 54 | *Df(3L)ED50002* | 22947 | 128631 | - |
| 23 | *Df(3L)BSC362* | 306168 | 628171 | - |
| 59 | *Df(3L)Exel6085* | 548528 | 749210 | - |
| 1<sup>♦</sup> | *Df(3L)ED4196* | 639583 | 1478937 | ++ |
| 27 | *Df(3L)BSC289* | 1332329 | 1628100 | - |
| 41 | *Df(3L)BSC800* | 1628101 | 1647451 | - |
| 46 | *Df(3L)BSC181* | 1688724 | 1841694 | - |
| 64 | *Df(3L)Aprt-32* | 1795318 | 2555775 | + |
| 45 | *Df(3L)ED4287* | 1795442 | 2551761 | - |
| 22 | *Df(3L)BSC119* | 2600282 | 2823614 | - |
| 35 | *Df(3L)Exel6092* | 2821245 | 3047162 | - |
| 32 | *Df(3L)BSC671* | 2982129 | 3193143 | - |
| 14 | *Df(3L)BSC672* | 3081311 | 3206906 | - |
| 37 | *Df(3L)ED4293* | 3226338 | 3250564 | - |
| 60 | *Df(3L)BSC368* | 3759821 | 4040635 | - |
| 55 | *Df(3L)BSC884* | 5601375 | 5770185 | - |
| 34 | *Df(3L)BSC410* | 5763773 | 6483285 | - |
| 39 | *Df(3L)BSC411* | 5969060 | 6618726 | - |
| 19 | *Df(3L)Exel6109* | 6736213 | 6936639 | - |
| 53 | *Df(3L)BSC27* | 6936605 | 7136086 | - |
| 58 | *Df(3L)BSC224* | 6957557 | 7150109 | - |
| 50 | *Df(3L)BSC33* | 7242439 | 7350373 | - |
| 21 | *Df(3L)BSC117* | 7242575 | 7328086 | - |
| 30 | *Df(3L)Exel8104* | 7353086 | 7522363 | - |
| 13 | *Df(3L)BSC375* | 7510880 | 7904179 | + |
| 12 | *Df(3L)BSC388* | 7643513 | 8184286 | - |
| 63 | *Df(3L)Exel6112* | 8089573 | 8351924 | - |
| 56 | *Df(3L)BSC815* | 8256164 | 8499740 | - |
| 11 | *Df(3L)BSC389* | 8415284 | 8582696 | + |
| 16 | *Df(3L)BSC816* | 8632181 | 8738462 | - |
| 24 | *Df(3L)ED4421* | 8738426 | 9377175 | - |
| 47 | *Df(3L)BSC113* | 9342609 | 9416591 | - |
| 65 | *Df(3L)AC1* | 9351951 | 10140553 | + |
| 17 | *Df(3L)BSC391* | 9446770 | 9697191 | - |
| 26 | *Df(3L)BSC118* | 9508772 | 9690291 | - |
| 44 | *Df(3L)BSC392* | 9671802 | 9892354 | - |
| 15* | *Df(3L)BSC673* | 9756714 | 10174058 | + |
| 48 | *Df(3L)BSC439* | 10507047 | 10964106 | - |
| 20 | *Df(3L)ED4470* | 11090089 | 11826330 | - |
| 49 | *Df(3L)ED4475* | 11580140 | 12401701 | - |
| 38 | *Df(3L)BSC730* | 12156077 | 12836424 | - |
| 25 | *Df(3L)BSC12* | 13037536 | 13221789 | - |
| 18 | *Df(3L)ED4543* | 13928325 | 14751140 | - |
| 33 | *Df(3L)BSC845* | 15504128 | 15819023 | - |
| 51 | *Df(3L)ED4606* | 16080584 | 16773223 | - |
| 6 | *Df(3L)ED4674* | 16654384 | 17042518 | + |
| 7 | *Df(3L)BSC414* | 16962973 | 17469226 | + |
| 62 | *Df(3L)Exel6132* | 17414682 | 17526191 | - |
| 57 | *Df(3L)ED4710* | 17487463 | 18139299 | - |
| 5 | *Df(3L)BSC775* | 17788244 | 18891426 | + |
| 43 | *Df(3L)BSC220* | 18965662 | 19164368 | - |
| 8* | *Df(3L)BSC839* | 20313247 | 20486308 | + |
| 10 | *Df(3L)BSC797* | 20445923 | 20942833 | - |
| 31 | *Df(3L)BSC449* | 20850015 | 21196030 | - |
| 40 | *Df(3L)BSC553* | 20984731 | 21219092 | - |
| 3 | *Df(3L)BSC419* | 21218032 | 21597878 | ++ |
| 2 | *Df(3L)BSC435* | 21304739 | 21770618 | ++ |
| 36 | *Df(3L)BSC836* | 21382499 | 21497772 | - |
| 52 | *Df(3L)BSC418* | 21382499 | 21637924 | - |
| 9 | *Df(3L)BSC223* | 21909520 | 22078536 | - |
| 4<sup>a</sup> | *Df(3L)BSC451* | 22069194 | 22684788 | ++ |
| 42 | *Df(3L)ED230* | 22127751 | 22827471 | - |
| 29 | *Df(3L)ED5017* | 22828597 | 22991401 | - |
| 61 | *Df(3L)1-16* | 24292305 | 24536634 | - |
| 28 | *Df(3L)6B-29* | 24977118 | 25115180 | - |
| <sup>a</sup> ♦ suggested haploinsufficiency<br><sup>b</sup> Breakpoints are as determined by the Bloomington stock center.<br><sup>c</sup> +, p value < 0.05; ++, p value < 0.01. P-values obtained from a Chi-square test. |  |  |  |  |

### Histone *H3* gene copy number is a modifier of *Pc* function

Nine annotated genes reside within the defined genomic interval: *CG12971, CG32436, CG32437, CG32440, ebd2, Pc, Rab26, Tbc1d8-9,* and *Tsr1* (**Figure 4B**). To determine which of these genes contributes to viability of flies with reduced histone gene copy number, we generated *H3.3A^Δ^; 12xHWT* animals heterozygous for single gene mutations. *CG12971*, *CG32437,* and *CG32440* were not tested because no loss-of-function alleles exist. Heterozygous MiMIC transposon insertion alleles of *ebd2, Rab26, Tbc1d8-9, Tsr1*, and *CG32436* did not impact viability of *H3.3A^Δ^; 12xHWT* flies (**Figure 4C**). However, three independent alleles of the *Polycomb* gene—*Pc^1^*, *Pc^3^* and *Pc^15^*—resulted in significant reductions in viability of *H3.3A^Δ^; 12xHWT* flies (55.0%, 61.8% and 52.4% of expected survive to adulthood, respectively) (**Figure 4C**). These data suggest that *H3.3A^Δ^; 12xHWT* flies are less viable when Polycomb function is reduced.

Polycomb group genes encode evolutionarily conserved regulators of cell identity. Polycomb complexes function to heritably silence expression of master regulator genes, including the Hox genes, which specify segmental identity (Kassis et al. 2017). In adult males, reduction of Polycomb function can result in homeotic transformations whereby the second (T2) and third (T3) thoracic legs acquire morphological features normally found only on the first thoracic legs (T1). This is most notably manifest by the appearance of sex combs on T2 and T3 legs, which normally only occur on T1 legs (Kaufman et al. 1980; Pattatucci et al. 1991). Based on our identification of the *Pc* gene in our genetic screen, we hypothesized that reduced histone gene copy number compromises Polycomb complex function. A prediction of this hypothesis is that reduced histone gene copy number would enhance *Polycomb* mutant phenotypes. Therefore, we evaluated the frequency and severity of homeotic transformations in *H3.3A^Δ^; 12xHWT*; *Pc/+* animals. We observed that *H3.3A^Δ^; 12xHWT* males heterozygous for a *Pc* null mutation (H3.3A*^Δ^; 12xHWT; Pc^3/+^)* exhibit an increased frequency of ectopic sex combs (100%) on T2 and T3 legs relative to *Pc^3/+^* (48%) or *H3.3A^Δ^; 12xHWT* males (0%) (**Figure 5A, Table 3**). Moreover, the expressivity of the ectopic sex comb phenotype is more severe in *H3.3A^Δ^; 12xHWT; Pc^3/+^* animals relative to *Pc^3/+^* controls, often having a full set of sex combs on T2 and T3 legs. Males and females of these genotypes also exhibit defects in posterior wing morphology, suggesting partial wing to haltere transformation due to a failure to maintain proper repression of *Ubx* in the wing (**Figure 5C**). Consistent with this hypothesis, immunostaining of *H3.3A^Δ^; 12xHWT; Pc^15/+^* third instar imaginal wing discs revealed ectopic *Ubx* expression in the pouch region (**Figure 5B**). We conclude that histone *H3* gene copy number contributes to Polycomb function during development.

**Figure 5.**
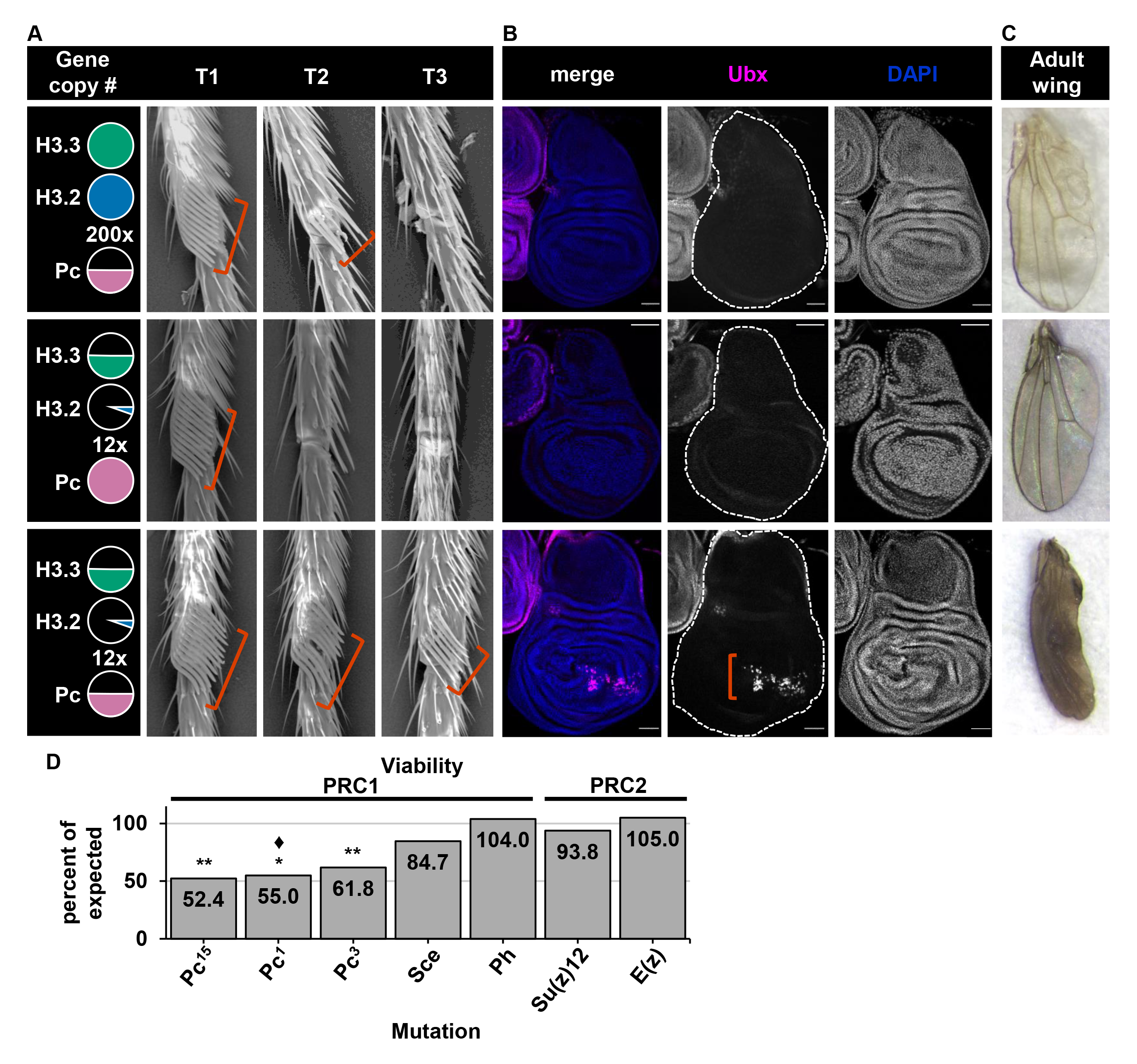
Reduced histone gene copy number disrupts *Polycomb* function. (A) Scanning electron micrographs of first (T1), second (T2), and third (T3) thoracic legs from adult males of the indicated genotypes, depicted as in Figure 1 with *Pc* gene dose in pink. Red brackets indicate the location where sex combs developed. (B) Confocal images of wing imaginal discs of the indicated genotypes stained for DAPI (blue) and Ubx (magenta). Red brackets indicate the wing pouch where ectopic Ubx expression occurred. Bars, 50 μM. (C) Bright field images of adult wings from the indicated genotypes. (D) Bar plots of viability of *H3.3A^Δ^; 12xHWT* animals heterozygous for mutations of PRC1 and PRC2 Polycomb complex members. Data are plotted as in Figure 3. Asterisks indicate statistical significance by Chi-square test (** p value < 0.01, ♦ potential haploinsufficiency irrespective of *H3* gene copy number, see Figure 5 in File S1 for p values).

**Table 3.**
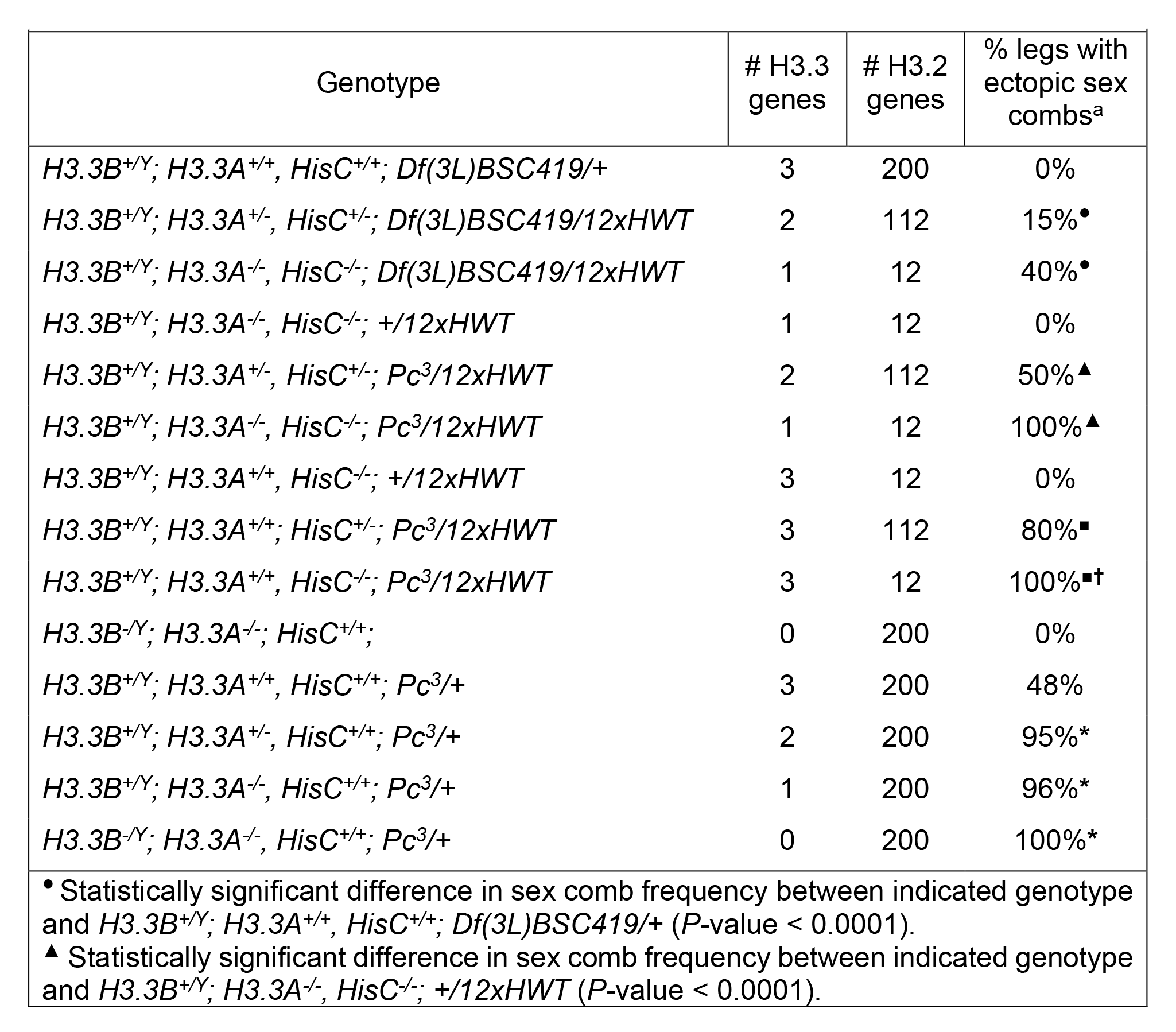
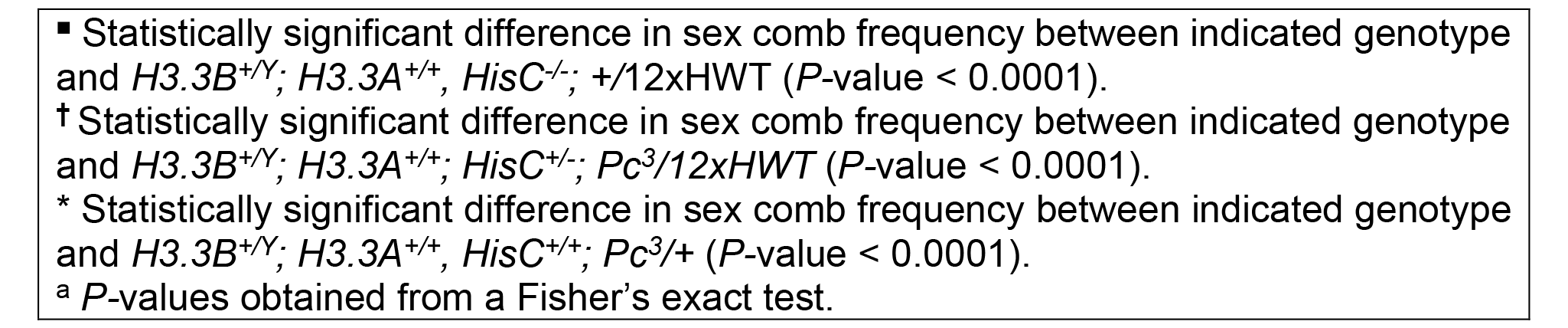
Decrease in *H3.2* and *H3.3* gene copy number increases frequency of ectopic sex combs.

Next, we determined whether mutations in other Polycomb group genes cause effects similar to mutations of *Pc* when histone *H3* gene copy number is reduced. *H3.3A^Δ^; 12xHWT* flies heterozygous for null mutations in *Sce* and *Ph*, which encode members of Polycomb Repressive Complex 1 (PRC1), are viable at expected frequencies (**Figure 5D**). Similarly, heterozygous mutations in Polycomb Repressive Complex 2 (PRC2) genes, *E(z)* and *Su(z)12*, do not cause reductions in viability (**Figure 5D**). *Pc* is a core component of PRC1, suggesting that the function of this Polycomb complex component is particularly sensitive to histone *H3.2* and *H3.3* gene copy number.

### Histone *H3.2* and *H3.3* gene copy number are each critical for Polycomb-mediated gene silencing

Since both *H3.2* and *H3.3* gene copy number are reduced in *H3.3A^Δ^; 12xHWT* animals, we next determined the individual requirement of either *H3.2* or *H3.3* gene copy number in Polycomb-mediated silencing. We quantified viability and assessed whether Polycomb target genes were de-repressed in either *12xHWT* or *H3.3^Δ^* animals that were also heterozygous for a *Pc* null mutation (*12xHWT; Pc^3/+^ and H3.3^Δ^; Pc^3/+^*, respectively). *12xHWT; Pc^3/+^* animals are viable at expected frequencies but have an increased frequency of ectopic sex combs on T2 and T3 legs like *H3.3A^Δ^; Pc^3/+^; 12xHWT* animals. *12xHWT; Pc^3/+^* animals also exhibit ectopic *Ubx* expression in the wing pouch of third instar imaginal discs and defects in adult posterior wing morphology (**Table 3**, **Figure 6A- D**). *112xHWT; Pc^3^*/+ animals are viable at the expected frequency but also exhibit increased frequencies of ectopic sex combs on T2 and T3 legs relative to *Pc^3/+^* controls (**Table 3**). Of note, the frequency of ectopic sex combs in *112xHWT; Pc^3/+^* animals (80%) is significantly lower than *12xHWT; Pc^3/+^* animals (100%) (**Table 3**). Thus, reducing *H3.2* gene dose makes animals sensitive to reduced Polycomb function, and the severity of homeotic transformation is proportional to canonical histone gene copy number. We also found that normal *H3.3* gene dose is necessary for Polycomb function. *H3.3^Δ^; Pc^3/+^* animals are not fully viable, with only 59.6% of expected surviving to adulthood (**Figure 6A**). *H3.3^Δ^; Pc^3/+^* males also exhibit increased frequencies of ectopic sex combs on T2 and T3 legs (100%), defects in adult posterior wing morphology, and *Ubx* de-repression in the wing pouch of third instar imaginal wing discs (**Figure 6B-D, Table 3**). Animals with one or two copies of *H3.3* survive to adulthood at a similar frequency—79.7% and 77.6% of expected, respectively—and animals with three copies of *H3.3* are viable at the expected frequency (**Figure 6A**). Consistent with these observations, males with only one copy of *H3.3B* also exhibit ectopic sex combs on T2 and T3 legs (**Table 3**). Taken together, these findings indicate that both *H3.2* and *H3.3* gene copy number are independently important for Polycomb-mediated epigenetic silencing, but viability is only affected by reduction in *H3.3* gene copy number.

**Figure 6.**
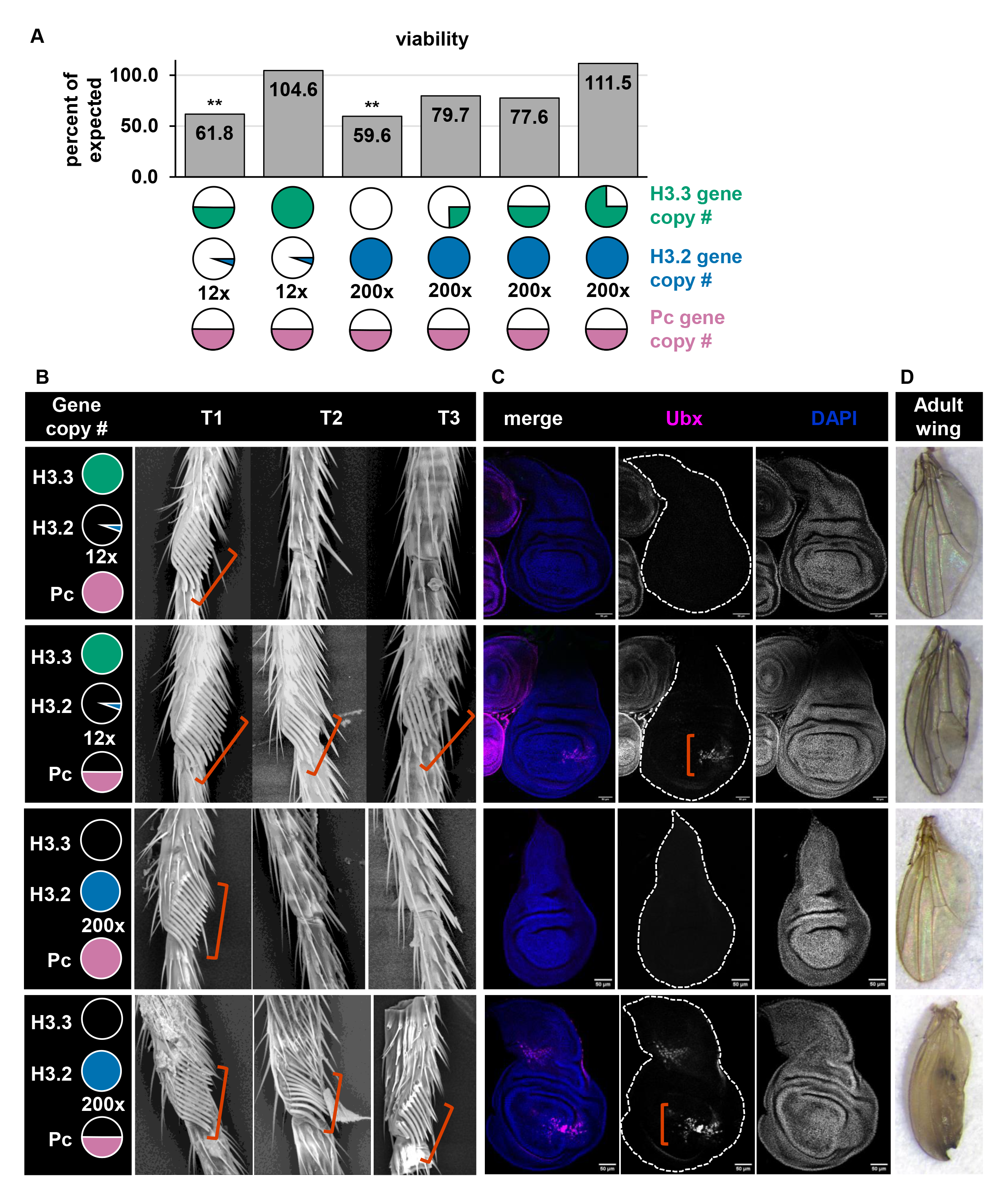
Polycomb function is sensitive to both *H3.2* and *H3.3* gene copy number. (A) Bar plot of viability for the indicated genotypes, depicted as in Figure 1 with *Pc* gene dose in pink. Data are plotted as in Figure 3. Asterisks indicate statistical significance by Chi-square test (** p value < 0.01, see Figure 6 in File S1 for p values). (B) Scanning electron micrographs of first (T1), second (T2), and third (T3) thoracic legs from adult males of the indicated genotypes. Red brackets indicate the location where sex combs developed. (C) Confocal images of wing imaginal discs of the indicated genotypes stained for DAPI (blue) and Ubx (magenta). Red brackets indicate the wing pouch where ectopic Ubx expression occurred. Bars, 50 μM. (D) Bright field images of adult wings of the indicated genotypes.

## Discussion

In this study we found that reducing either canonical or variant histone *H3* gene copy number disrupts Polycomb-mediated gene repression. Two major protein complexes establish and maintain Polycomb-mediated repression: Polycomb Repressive Complex 1 and 2 (PRC1 and PRC2). PRC2 catalyzes H3K27me3, both PRC2 and PRC1 bind to H3K27me3, and PRC1 facilitates repression of local chromatin (Blackledge and Klose 2021). Mutations in core components of PRC1 and PRC2 disrupt the formation of these domains and cause transcriptional de-repression of Polycomb targets, such as Hox genes (Kennison and Tamkunt 1988; Paro 1990; Orlando 2003). We found that reduction in canonical or variant *H3* gene copy number results in homeotic transformations associated with de-repression of the Polycomb target genes *Ubx* (posterior wing transformation) and *Scr* (ectopic sex comb development in males). Consistent with these findings, a previous study in *D. melanogaster* found that heterozygosity of *HisC* suppresses homeotic transformation phenotypes in animals with a mutation that causes ectopic silencing of Polycomb target genes (Bajusz et al. 2001). Reduction of canonical histone gene copy number in *D. melanogaster* also modifies position-effect variegation, a phenomenon mediated by H3K9me3-marked constitutive heterochromatin (Moore et al. 1979; Moore et al. 1983; Sinclair et al. 1983). Moreover, deletion of all variant *H3.3* gene copies in mouse embryonic fibroblasts results in disruption of heterochromatin domains at pericentromeric repeat regions, centromeres, and telomeres (Jang et al. 2015). Collectively, these findings indicate that *H3.2* and *H3.3* gene copy number play an important role in establishing efficient silencing via H3K9me3- and H3K27me3-mediated heterochromatin. We discuss below potential mechanisms for how changes in histone gene copy number might impact Polycomb-mediated repressive chromatin.

### Histone protein abundance and stoichiometry may influence Polycomb-mediated repressive chromatin

Here we show that although *12xHWT* animals develop normally, reducing *Pc* gene dose by half in a *12xHWT* background results in mutant phenotypes associated with impaired Polycomb-mediated gene silencing. This genetic interaction suggests that the combination of reduced amounts of canonical histones and Polycomb prevents the proper formation of a repressive chromatin domain at Polycomb-silenced genes. However, our previous work found that canonical histone transcript levels are similar in *12xHWT* and *200xWT* animals, at least in early embryos (11), and here we found similar levels of total H3.3 and H3 protein in *12xHWT* and *200xWT* animals by western blotting. H3.2 and H3.3 are highly abundant proteins, and western blots may not provide the sensitivity needed to identify small changes in H3 protein levels that could be biologically meaningful. Moreover, a subtle decrease in histone protein abundance may result in changes in nucleosome occupancy that preferentially affect heterochromatin function. Polycomb chromatin domains have elevated nucleosome occupancy and decreased nucleosomal spacing and therefore may be particularly sensitive to changes in histone abundance (King et al. 2018). In fact, disruption of PRC1-mediated chromatin compaction in *D. melanogaster* results in de-repression of Hox genes (Bonnet et al. 2022). In addition to direct effects of decreased histone abundance at Polycomb target genes, it is also possible that indirect effects contribute to Polycomb target gene misregulation in *12xHWT* animals. Previous work showed that reductions in the concentration of free histone H3 results in increased local histone recycling during replication in *Xenopus* egg extracts and *D. melanogaster* embryogenesis (Gruszka et al. 2020; Mühlen et al. 2023b). Thus, another possibility is that reduced histone gene copy number results in an increased proportion of recycled histones within chromatin. If recycled histones carry PTMs that antagonize Polycomb function, such as H3K36me3, they could alter the PTM landscape and impact target gene repression at Polycomb domains (Finogenova et al. 2020; Bonnet et al. 2022; Mühlen et al. 2023b; Salzler et al. 2023). Future studies examining chromatin accessibility and the PTM landscape at Polycomb target domains upon reduction in *H3* gene copy number would help address these issues.

Nucleosomes and histone-chaperone complexes are multiprotein complexes that assemble with defined stoichiometries (Luger et al. 1997; Andrews and Luger 2011; Grover et al. 2018), and many genomic processes are sensitive to perturbations in subunit stoichiometry within these complexes. For instance, disrupting the stoichiometric balance between H2A:H2B dimers and H3:H4 dimers in yeast causes genome instability and mitotic chromosome loss (Meeks-Wagner and Hartwell 1986). In all of the genotypes we assessed that display mutant phenotypes indicative of impaired Polycomb repression, the balance between *H3.2* and *H3.3* gene copy number is altered, and this change in the relative abundance of H3.2 and H3.3 could impact genome regulation. Consistent with this interpretation, work done in mice shows that displacement of H3.3—and enrichment of replication dependent H3.1—at regulatory regions causes transcriptional deregulation and chromosomal aberrations (Chen et al. 2020). Thus, the stoichiometric balance between H3.2 and H3.3 in chromatin may be critical for maintaining Polycomb target gene silencing in flies.

A corollary to this model is that proper stoichiometric balance between H3 proteins and their chaperones is needed for repression of Polycomb targets. Previous work in mouse cells shows that H3.3-specific chaperones interact with PRC1 and PRC2, and that these interactions are needed for the recruitment of H3.3 to H3K9me3-dependent heterochromatin and for the establishment of H3K27me3 at developmental gene promoters (Banaszynski et al. 2013; Liu et al. 2020). Therefore, one could posit that altering the stoichiometric balance between H3.3 and its chaperones may perturb the establishment or maintenance of Polycomb domains. Notably, in our genetic screen for viability we identified a deficiency that covers *Yem*, an H3.3-specific chaperone, suggesting that mechanisms regulating the levels of canonical and variant histone within the genome involve control of histone deposition into chromatin.

### Distinct roles of canonical H3.2 and variant H3.3 in Polycomb-mediated silencing

Canonical *H3.2* and variant *H3.3* differ in their expression patterns and protein sequence. Our genetic analyses demonstrate that reducing *H3.3* gene copy number, but not *H3.2* gene copy number, causes a decrease in viability of *Pc* heterozygotes, suggesting *H3.2* and *H3.3* may have non-identical roles in Polycomb target gene regulation. *H3.3^Δ^* mutants do not uncouple *H3.2* expression from S phase, and *12xHWT* mutants still express *H3.3* throughout all of interphase. Thus, depleting the pool of H3 outside of S phase in the *H3.3^Δ^* mutants may sensitize cells to small perturbations in Polycomb-mediated silencing during development. For example, Polycomb Response Elements (PREs), such as those that regulate *Ubx,* are sites of high histone turnover even though they reside within silent chromatin domains (Mito et al. 2007). As such, PREs may be particularly sensitive to the loss of available H3 protein outside of S phase. Reduced histone occupancy at PREs could impact Polycomb repression and organismal viability. Our finding that viability of animals with reduced *H3.2* gene copies increases as *H3.3* gene copy number increases supports this model. Alternatively, canonical H3.2 and variant H3.3 proteins may have distinct functions at PREs or Polycomb target domains. H3.2 and H3.3 differ at residue 31 on the N-terminal tail and H3.3S31 can be phosphorylated (H3.3S31ph). It is known that histone H3 post-translational modifications can influence one another (Yuan et al. 2011; Finogenova et al. 2020). Notably, H3.3S31ph affects the local PTM landscape and binding of factors that interact with the H3 tail, like the H3K36me3 reader ZMYND11 (Armache et al. 2020; Sitbon et al. 2020). Conceivably, these H3.3S31ph-specific effects on the PTM landscape could impact Polycomb function. Future studies probing the impacts of an H3.3S31A mutation, which renders the residue non-modifiable, on Polycomb function would address this possibility.

In summary, our data investigating the control of canonical and variant histone abundance provide evidence that Polycomb-mediated gene silencing is sensitive to both canonical and variant histone gene copy number. This work advances our understanding of the distinct and overlapping functions of canonical and variant histones.

## Materials & Methods

### Fly Stocks & Husbandry

Fly stocks were maintained on standard corn medium provided by Archon Scientific (Durham, NC). See stocks in File S1 for a list of all stocks.

### CRISPR-mediated generation of *H3.3A^Δ^*

pCFD4 plasmids encoding dual gRNAs (gRNA_1: caaggcgccccgcaagcagc, gRNA_2: tgcaccgtgactatttcata) targeting *H3.3A* were injected into embryos expressing Cas9 from the *nanos* promoter (*y1 M{nos-Cas9.P}ZH- 2A w\**; RRID: BDSC_54590) by *GenetiVision* Corporation (Houston, TX). *H3.3A*^Δ^ alleles were identified by PCR of genomic DNA and confirmed by sequencing, which revealed a 265 bp deletion removing amino acids 19 through 94 of the *H3.3* open reading frame. The deletion breakpoints are indicated by “…/…” in the following sequence: CAAGGCGCCCCGCA…/…TACGGTCATGTAAT. The *H3.3A^Δ^* allele was determined to be amorphic because *H3.3B^0^; H3.3A^Δ^; 12xHWT/+* animals are inviable. CRISPR diagnostic screen primer set: H3.3A**^Δ^** for-CCCGATGAATATAGGGTCACAC, H3.3A**^Δ^** rev-CTGGATGTCCTTGGGCATAAT. pCFD4-U6:1_U6:3 was a gift from Simon Bullock (Addgene plasmid #49411; http://n2t.net/addgene:49411; RRID:Addgene_49411). *His3.3A* reference sequence: NCBI Gene ID 33736.

### Viability

To examine the effect of *H3.2* gene copy number on *H3.3^null^* viability (Figure 1D), the following four crosses were performed:

1. *H3.3B^0^ / H3.3B^0^; H3.3A^2×1^, ΔHisC, twistGal4 / CyO; x yw / Y; H3.3^A2×1^, ΔHisC, UAS-YFP / CyO; 12xHWT / 12xHWT*
2. *H3.3B^0^ / H3.3B^0^; H3.3A^2×1^, ΔHisC, twistGal4 / CyO; x yw / Y; H3.3^A2×1^, ΔHisC, UAS-YFP / CyO; 12xHWT-VK33, 8xHWT-86F6 / +*
3. *H3.3B^0^ / H3.3B^0^; H3.3A^2×1^, ΔHisC, twistGal4 / CyO; x H3.3B^0^ / Y; H3.3A^2×1^ / CyO, twistGFP*
4. *H3.3B^0^ / H3.3B^0^; H3.3A^Δ^ / CyO, twistGFP; x H3.3B^0^ / Y; H3.3A^Δ^ / CyO, twistGFP;*

To examine the effect of *H3.3* gene copy number on *12xHWT* viability (Figure 1D), the following 3 crosses were performed:

1. *H3.3B^0^ / H3.3B^0^; H3.3A^2×1^, ΔHisC, UAS-YFP / CyO; 12xHWT / 12xHWT x yw; H3.3^A2×1^, Df(2L)ΔHisC^Cadillac^ / CyO;*
2. *yw / yw; H3.3A^2×1^, Df(2L)Δ HisC^Cadillac^; 12xHWT / 12xHWT x H3.3B^0^ / Y; H3.3A^2×1^, ΔHisC, twistGal4 / CyO;*
3. *yw / yw; ΔHisC, twistGal4 / CyO; x yw; ΔHisC, UAS-YFP / CyO; 12xHWT / 12xHWT*

Vials were maintained at 25°C and flipped every other day. Data were plotted as percent observed of expected based on Mendelian ratios. Chi-squared analysis was performed to determine statistical significance. A significance threshold of p < 0.05 was used in this study. See stocks in File S1 for all genotypes and progeny numbers.

All genotypes were confirmed by PCR using the following primer sets:

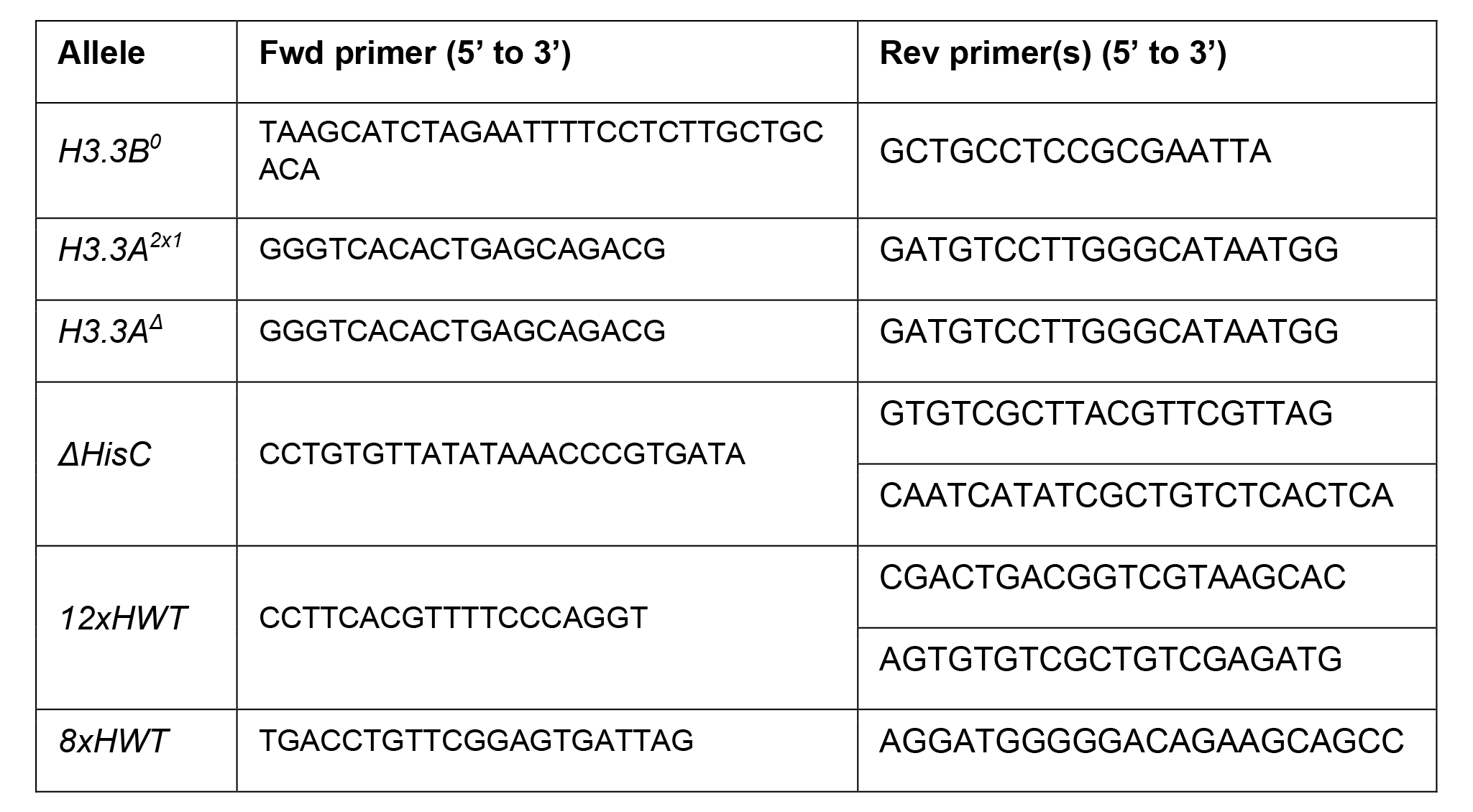

### RNA sequencing

For each replicate sample, 25 brains were dissected from wandering 3^rd^ instar larvae and homogenized in Trizol solution. RNA was isolated from the Trizol aqueous phase using the Zymo RNA Clean and Concentrator-5 kit (Genesee Scientific #11-352) including treatment with DNAse I, as per the manufacturer’s instructions. Libraries were prepared from polyA selected RNA using the KAPA stranded mRNA kit (Roche # 07962207001) and sequenced using the NOVASeq-S1 paired-end 100 platform. Sequence reads were trimmed for adaptor sequence/low-quality sequence using BBDuk (bbmap v38.67) with parameters: ktrim=r, k=23, rcomp=t, tbo=t, tpe=t, hdist=1, mink=11. Dm6 genome files for use with the STAR aligner were generated using parameters: sjdbOverhang 99. Paired-end sequencing reads were aligned using STAR v2.7.7a with default parameters (Dobin et al. 2013). featureCounts (subread v2.0.1) was used with default parameters to count reads mapping to features (Liao et al. 2014). DESeq2 (v1.34.0) was used to identify differentially expressed genes (Love et al. 2014). Differentially expressed genes were defined as genes with an adjusted P-value less than 0.05 and an absolute log2 fold change greater than 1.

### Western blots

Protein extracts from *H3.3^null^* third instar larvae and *12xHWT* and *yw* (*200xWT*) third instar larval wing discs were prepared by boiling samples for 10 minutes in Laemmli SDS-PAGE loading buffer followed by sonication using the Bioruptor Pico sonication system (Diagenode) for 10 cycles (30 sec on, 30 sec off). Samples were clarified by centrifugation. Proteins were fractionated on BioRad Any kD^TM^ Mini-PROTEAN^®^ TGX^TM^ Precast Protein Gels GX (BioRad #4569033) and were transferred to 0.2um nitrocellulose membranes (BioRad #1620112) at 100V for 10 minutes and 60V for 20 minutes. Total protein was detected using G-Bioscience Swift Membrane Stain^TM^ (G- Bioscience, 786677). Membranes were probed using the following antibodies: rabbit anti-H3 (1:60,000; Abcam Cat# ab1791, RRID:AB_302613), rabbit anti-H3.3 (1:1,000, Abcam Cat# ab176840, RRID:AB_2715502), and mouse anti-tubulin (1:15,000, Sigma-Aldrich Cat# T6074, RRID:AB_477582). Western blot analysis was performed using the following HRP conjugated secondary antibodies: goat anti-Mouse-IgG-HRP (1:10,000, Thermo Fisher Scientific Cat# 31430, RRID:AB_228307), donkey anti-Rabbit-IgG-HRP (1:10,000, (Cytiva Cat# NA934, RRID:AB_772206). Blots were detected using Amersham ECL Prime Western blotting Detection Reagent (Cytvia, RPN2232). ImageLab densitometry analysis was used to determine total protein, tubulin, H3.3, and H3 band intensity. Histone signal was normalized to corresponding tubulin signal. Normalized signals from different titrations of the same genotype were averaged and resulting values were set relative to the wild-type value. This process was completed for three biological replicates. See Figure in File S1 for raw data for each replicate.

### EdU + RNA FISH

Third instar larval eye discs were dissected in Grace’s medium and incubated in 0.1 mg/mL EdU for 30 minutes. Samples were then washed in phosphate-buffered saline (PBS) for 5 minutes, fixed for 30 minutes in 4% paraformaldehyde (16% paraformaldehyde, diluted in PBS), washed 3×15 minutes in PBS with Triton (PBST) (0.5% Triton X-100) and washed for 30 minutes in 3% Bovine serum albumin (BSA) in PBS. EdU was detected using the Click-iT EdU Cy5 imaging kit (Invitrogen) according to the manufacturer’s instructions. After EdU detection, samples were washed in 3% BSA for 15 minutes, washed in PBS for 5 minutes, and subsequently fixed in 4% paraformaldehyde for 15 minutes. Next, RNA-FISH was performed using Stellaris^®^ H3.2 mRNA probes with TAMRA fluorophores following the manufacturer’s instructions for *D. melanogaster* wing imaginal discs. Samples were stained with DAPI (1 ug/mL) at 37°C for 30 minutes in Wash Buffer A. The maximum projection from 4 adjacent z-slices from third instar wandering larval eye discs was used as representative images for each genotype.

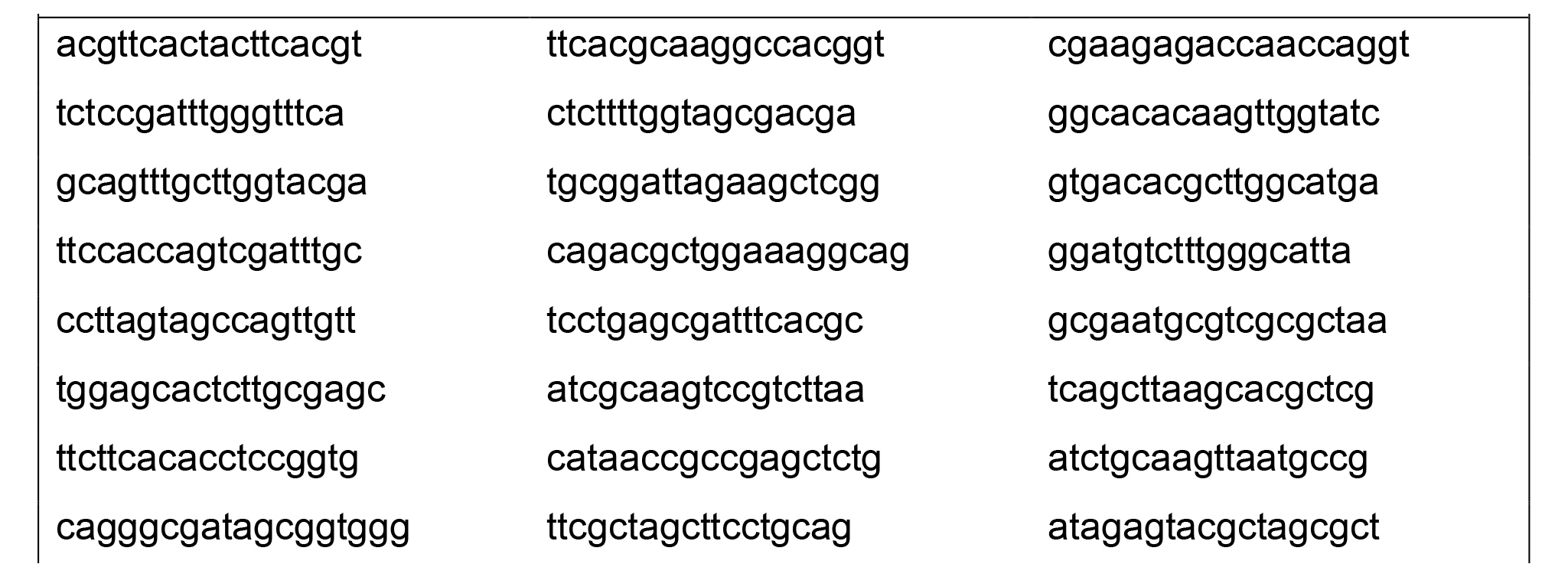
H3.2 RNA FISH probe set. Probes are oriented in the 3’ to 5’ direction.

### Genetic Screen Viability

Bloomington third chromosome deficiency stock males were crossed to *yw; H3.3A^2×1^, ΔHisC, twistGal4 / CyO; MKRS / TM6B* virgin females. Subsequently, *yw; H3.3A^2×1^, ΔHisC, twistGal4 / +; Df(3) / MKRS* male progeny were crossed with *yw; H3.3A^2×1^, HisC^Cadillac^ / H3.3A^2×1^, HisC^Cadillac^; 12xHWT / 12xHWT* virgin females. All fly stocks were maintained on standard corn medium at 25°C. Crosses were flipped every other day for 8 days. Progeny were scored once per day. Animals eclosing from deficiency crosses were counted beginning ten days post egg-laying based on the presence or absence of dsRed from the *HisC^Cadillac^* locus and stubble phenotype from MKRS until all adult flies eclosed. Expected and observed ratios of the desired genotypes were calculated following the completion of counting. Percent of expected was determined by the ratio of *yw; H3.3A^2×1^, ΔHisC, twistGal4 / H3.3A^2×1^, Cadillac; Df(3) / 12xHWT was* to *yw; H3.3A^2×1^, Cadillac / CyO; Df(3) / 12xHWT* siblings. Significance was determined by chi-square test; thresholds of p < 0.05 and p< 0.01 were used in this study. All genotypes and progeny numbers can be found in File S1.

### Immunofluorescence

For Ubx staining of wing discs, third instar larval cuticles were inverted and fixed in 4% paraformaldehyde in PBS for 20 minutes at room temperature. Cuticles were washed for 1 h in PBST (0.15% Triton X-100). Mouse anti-UBX (1:30, DSHB Cat# FP3.38, RRID:AB_10805300) was used overnight at 4°C. Goat anti-mouse IgG secondary antibody (1:1000, Thermo Fisher Scientific Cat# A-11029, lot #161153, RRID:AB_2534088) was used for 2 hours at room temperature. DNA was counterstained with DAPI (0.2ug/mL) and the discs were mounted in Vectashield^®^ (VWR, 101098-042) mounting media and imaged on a Leica Confocal SP8.

### Scanning electron microscopy

One-to four-day-old flies were dehydrated in ethanol and images of legs were taken using a Hitachi TM4000Plus tabletop SEM microscope at 15 kV and 500x magnification.

## Data Availability

Strains are available upon request. Raw RNA-seq data were deposited to Gene Expression Omnibus (GEO) under accession GSE228058. Additional information is available from the corresponding authors upon request.

## Acknowledgements

The authors thank Aaron Crain for the generation of the *Df(2L)ΔHisC^Cadillac^*allele.

